# *Drosophila* DNA/RNA methyltransferase contributes to robust host defense in ageing animals by regulating sphingolipid metabolism

**DOI:** 10.1101/362012

**Authors:** Varada Abhyankar, Bhagyashree Kaduskar, Siddhesh S. Kamat, Deepti Deobagkar, Girish Ratnaparkhi

**Affiliations:** Department of Zoology, Savitribai Phule Pune University, Pune 411007, INDIA; Department of Biology, Indian Institute of Science Education & Research, Pune 411008, INDIA; ISRO Chair Professor, Savitribai Phule Pune University, Pune 411007, INDIA

**Keywords:** DNA methyltransferase, *Drosophila*, Lipid homeostasis, hemocyte, innate immunity, ceramide, signaling, sphingosine-1-phosphate (S1P), Methyltransferase, S1P, hemocyte shape, lipid homeostasis, age dependence, robust immune response

## Abstract

*Drosophila* methyltransferase (*Mt2*) has been implicated in methylation of both DNA and tRNA. In this study, we demonstrate that loss of *Mt2* activity leads to an age dependent decline of immune function in the adult fly. A newly eclosed adult has mild immune defects that exacerbate in a fifteen-day old *Mt2*^−/−^ fly. The age dependent effects appear to be systemic, including disturbances in lipid metabolism, changes in cell shape of hemocytes and significant fold changes in levels of transcripts related to host defense. Lipid imbalance, as measured by quantitative lipidomics, correlates with immune dysfunction with high levels of immunomodulatory lipids, sphingosine-1phosphate (S1P) and ceramides, along with low levels of storage lipids. Activity assays on fly lysates confirm the age dependent increase in S1P and concomitant reduction of S1P lyase activity. We hypothesize that *Mt2* functions to regulate genetic loci such as S1P lyase and this regulation is essential for robust host defense as the animal ages. Our study uncovers novel links between age dependent *Mt2* function, innate immune response and lipid homeostasis.

## INTRODUCTION

Innate immunity (Janeway and Medzhitov, 2002) is an evolutionary conserved host defense mechanism present throughout the plant and animal kingdoms. It is the predominant form of defense against pathogens. Invertebrates lack the adaptive immune system and are thus an excellent model to study innate defense mechanisms in isolation. The fruit fly, *Drosophila melanogaster*, responds to microbial infections by mounting a defense (Akira et al., 2006; Anderson, 2000; Buchon et al., 2014; Ferrandon et al., 2007; Iwasaki and Medzhitov, 2010; Lemaitre and Hoffmann, 2007; Ligoxygakis, 2013; Uvell and Engstrom, 2007) against the invading organisms. The first line of defense is the external cuticle and epithelial barriers. Once the pathogen breaches these barriers and reaches the hemocoel, they encounter systemic defenses, both humoral and cellular. The humoral response encompasses the up-regulation of the defense genes and antimicrobial peptides (AMPs) from the fat body of *Drosophila*, melanization and the release of reactive oxygen species, while hemocytes (Agaisse et al., 2003; Williams, 2007) lead the cellular response, by efficiently phagocytosing and encapsulating microorganisms. Defense genes thus encode proteins/RNA that function to counteract the effect of the invader and repair the damage caused. The regulation and thereby expression of defense genes is controlled by a number of well-characterized signal transduction pathways like the Toll signaling pathway, Immune-deficient (IMD) pathway, c-Jun N-terminal kinases (JNK) and the JAK-STAT pathways (Agaisse et al., 2003; Delaney et al., 2006; Govind and Nehm, 2004; Kounatidis and Ligoxygakis, 2012; Lemaitre and Hoffmann, 2007; Lemaitre et al., 1996; Matova and Anderson, 2010; Schneider, 2007; Silverman et al., 2003). Extracellular ligands and/or cell surface receptors sense signatures of systemic microorganisms and this signal is transduced via the aforementioned transduction pathways to activate the *Drosophila* NF**κ**B’S Dorsal, Dif and Relish (Brennan and Anderson, 2004; Govind, 1999; Hetru and Hoffmann, 2009; Ip et al., 1993; Kounatidis et al., 2017; Lemaitre et al., 1995; Tanji and Ip, 2005).

The immune response in *Drosophila* shows complex age-dependent phenotypes (Clark et al., 2014; Zerofsky et al., 2005). In terms of the cellular response, phagocytic activity declines by 30% in one month old flies and this correlates with a decline in number of hemocytes (Horn et al., 2014; Mackenzie et al., 2011). Levels of expression of many defense genes vary greatly with age (Felix et al., 2012; Zerofsky et al., 2005), suggesting age dependent regulation of the immune response. The overall picture is complex and suggests compensatory mechanisms to deal with infection while ageing. Longevity has also been linked to immune function with many critical signaling networks that regulate longevity such as Insulin-IGF like (IIL) and TOR pathways (Grewal, 2009; Johnson et al., 2013; Kapahi et al., 2017; Partridge et al., 2011) shown to communicate with the central immune pathways for robust regulation of host defense (DeVeale et al., 2004; Kounatidis et al., 2017; Unckless et al., 2015).

In this study, we characterize the immune response in flies with perturbation in activity of *Drosophila DnMt2* (called *Mt2* henceforth), a cryptic DNA/RNA methyltransferase (MT). Vertebrates have multiple DNA MTs, classified as *DnMt1*, *Mt2*, *DnMt3a*, *DnMT3b*, based on their activity and structural features (Basu et al., 2016; Okano et al., 1998). In contrast, Mt2 is the only MT identified in *Drosophila* (Tang et al., 2003). Originally, Mt2 was characterized as a DNA-MT, but recent research suggests that Mt2 might function primarily as a RNA-MT (Goll et al., 2006; Schaefer et al., 2010), with methylation enhancing tRNA stability. *Mt2* null flies (*Mt2*^−/−^) do not show overt developmental abnormalities and their lifespan is near normal under non-stressed conditions. Under stress (Becker et al., 2012; Schaefer et al., 2010; Thiagarajan et al., 2011), *Mt2*^−/−^ flies show a shorter lifespan (Lin et al., 2005). Flies grown in overcrowded conditions develop melanotic spots (Durdevic et al., 2013), suggesting disturbances in immune function. Infection studies also suggest that Mt2 plays an important role in acute immune response to *Drosophila* C virus (DCV) by binding to and possibly methylating viral RNA (Durdevic et al., 2013).

Here, we demonstrate that *Mt2*^−/−^ flies show an age dependent immune decline. The ability of adult flies to clear bacteria decreases dramatically by the fifteenth day post eclosion. Adult hemocytes are sickle-shaped with numbers in excess of that for a wild type animal of the same age. The age dependent effects are correlated with perturbations in lipid homeostasis, suggesting that the decline may be a direct response to changes in critical lipid molecules involved in cellular homeostasis. We hypothesize that Mt2 regulates enzymes involved in lipid homeostasis and this function is essential for supporting a robust immune response as the animal ages.

## RESULTS

### Mt2-Null flies show reduction in life-span after bacterial infection

Earlier reports on *Mt2*^−/−^ flies indicated that these flies show a shortened lifespan (Lin et al., 2005), are sensitive to stress (Schaefer et al., 2010) and are susceptible to viral infection (Durdevic et al., 2013). In our study, we first confirmed that *Mt2*^−/−^ flies had shorter lifespan (Fig. 1A) and then tested if *E. coli.* infection had an effect on *Mt2*^−/−^ lifespan. *Mt2*^+/+^, *Mt2*^−/−^ and *Mt2*-*(Transgenic Rescue)TG* lines (genotypes as described in Materials & Methods) were either infected with *E. coli* or mock infected with sterile 1X PBS. Infection of *Mt2*^−/−^ flies, when compared to *Mt2*^+/+^, had an increased rate of lethality. In contrast, *Mt2*-*TG* animals showed a near normal lifespan for both mock and infection experiments, suggesting a role for Mt2 in host defense against gram negative bacteria. In order to get a more detailed picture for roles for Mt2 in the innate immune response, as described in following sections, we tested the functionality of both the cellular and humoral arms of the immune response by bacterial clearance assays as well by measuring change in transcript levels of defense genes in *Mt2*^−/−^ flies before and after infection.

**Fig. 1:**
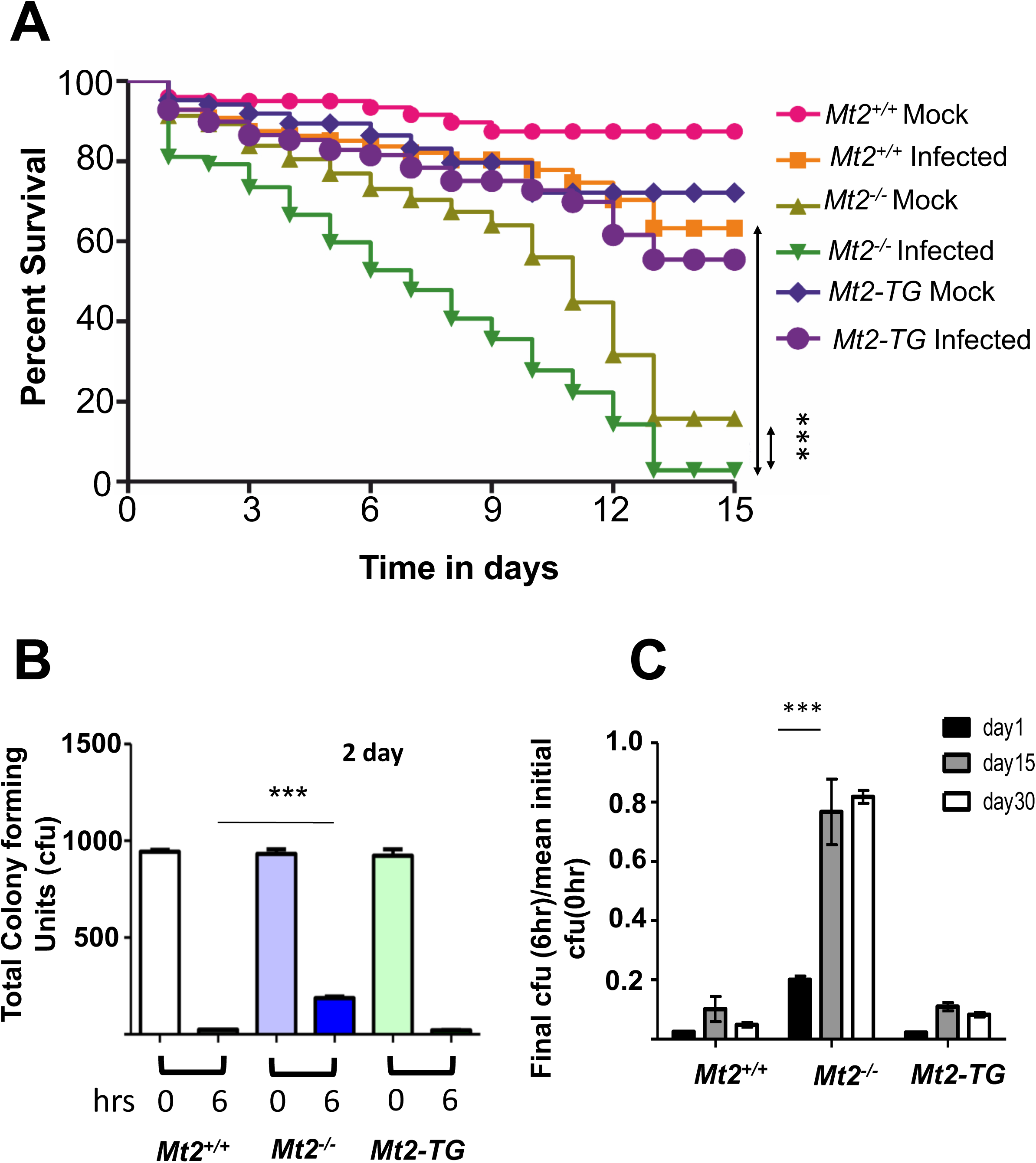
Life span and bacterial clearance assays suggest a decline in immune function for Mt2^−/−^ flies. A. Mock infected *Mt2*^−/−^ flies show a shorter lifespan when compared to mock infected *Mt2*^+/+^ animals. When infected with *E. coli*, *Mt2*^−/−^ flies show enhanced lethality at 25 °C and 29 °C (data not shown), when compared to infected *Mt2*^+/+^ flies. Infected *Mt2*^−/−^ flies show increased mortality as compared to mock infected *Mt2*^−/−^ flies. Experiments were in biological triplicates, with life spans curves analyzed using Log-rank (Mantel-Cox) Test, in GraphPad Prism version 5. Analysis indicates that the curves *Mt2*^−/−^ (mock) vs *Mt2*^−/−^ (infected) as well as the *Mt2*^−/−^ (infected) vs *Mt2*^+/+^ (infected) differ significantly (p<0.0001).
B. Total bacterial colony forming unit (cfu) count at 0h and 6h post infection for 2 day old *Mt2*^+/+^, *Mt2*^−/−^ and *Mt2*-*TG* males at 25 °C. The *Mt2*^−/−^ flies fail to clear bacterial load to the same extent as wild type, 6h post infection. The data represents three independent biological replicates. Data analyzed by 1-way ANOVA. For all figures henceforth, ‘^∗^’, indicates p<0.01, ‘^∗∗^’, p<0.05 and ‘^∗∗∗^’, p<0.001.
C. Cfu count for ageing flies at day 1, day 15 and day 30 post eclosion. The 6h post-infection cfu was normalized to the mean 0h cfu in each case. 2-way ANOVA post arcsine transformation was used to test significance. 1 day old *Mt2*^−/−^ flies showed a mild deficiency in their ability to clear bacteria, which worsened dramatically with age. N (biological replicates) =3, n (number of flies, for each day/time-point and each genotype) =4.

### *Mt2*-Null flies show age dependent impairment in bacterial clearance

We infected 2 day old adult *Mt2*^+/+^, *Mt2*^−/−^ and *Mt2*-*TG* with a saturated, Ampicillin (Amp) resistant culture of *E. coli.* Six hours post infection, the animals were crushed and processed, as described in Materials & Methods, to measure the decrease in *E. coli* numbers as a consequence of clearance by a robust immune response. The *Mt2*^−/−^ flies were an order of magnitude less efficient (Fig. 1B, compare 6 Hour *Mt2*^+/+^ with *Mt2*^−/−^) in clearing the infection as against the wild type or the rescue line (Fig. 1B). 2 day old *Mt2*^−/−^ flies thus, are impaired in their ability to clear bacteria, suggesting that Mt2 activity supports host defense against bacteria.

Next, we performed age dependent analysis for the ability of adult flies to clear infection. Surprisingly, we found that *Mt2*^−/−^ flies showed significant age-dependent loss in their ability to clear bacterial infection as compared to wild type flies; an ability regained by replacing *Mt2*, as in the *MT2*-*TG* flies (Fig. 1C). Wild type flies did not show significant loss in their ability to clear infections over 30 days. In stark contrast, 15 days *Mt2*^−/−^ flies cleared bacteria 8-10 fold less efficiently. 30 day old flies showed similar deficiency, suggesting that there is a steep decline in ability to clear infection from day 2 to day 15.

### *Mt2*-Null animals have age dependent defects in hematopoiesis

The earliest difference between *Mt2*^+/+^ and *Mt2*^−/−^ we could find in the cellular response was in the third instar larvae. We found that crystal cells, which are platelet like cells involved in melanization, are higher in number in *Mt2*^−/−^ animals as compared to *Mt2*^+/+^ (Suppl. Fig. 1A). This indicated that numbers of blood cells are not as well-regulated in the mutant. This data led us to look closely at the number of hemocytes in adults as they age (Fig. 2A). 15 day old wild type animal had fewer number of hemocytes, when compared to a 2 day old fly. In contrast, the number of hemocytes significantly increased in the *Mt2*^−/−^ with age, a trend opposite to that of the wild type fly and the *Mt2*-*TG* line. This would indicate that the increase in hemocytes with age is a *Mt2*^−/−^ specific event. While counting the hemocytes using light microscopy, we also noticed that the hemocytes in *Mt2*^−/−^ animals had ellipsoid, rice grain like shape as compared to circular shapes in the wild-type hemocytes. To get a clearer picture we employed Scanning electron microscopy (SEM; See *Materials and Methods)*. When compared to round wild type hemocytes, the Mt2^−/−^ hemocytes appeared flat, folded and C-shaped (Fig. 2B), reminiscent of diseased Sickle shaped human RBC’s. Quantitation of the roundness index of the image SEM data indicated a dramatic change in shape of the hemocytes in *Mt2*^−/−^ animals (Fig. 2B, C). This change in cell shape could account for the inefficiency of the *Mt2*^−/−^ hemocytes in clearing the bacterial load in the animal. Based on the above results, we tested transcript levels of *serpent (srp)*, a gene involved in regulating hemocyte morphology and phagocytic function (Petersen et al., 1999; Ramet et al., 2002; Shlyakhover et al., 2018) for 15 day old flies. *Srp* shows reduction in transcript levels (Fig. 2D). This suggests that Mt2 regulates *srp*/Srp expression directly or indirectly, affecting the cellular arm of immunity. We then measured transcript levels, for 15 day old animals for *eater* (Kroeger et al., 2012) and *u*-*shaped* (Muratoglu et al., 2007), genes known to be critical for hemocyte phagocytosis and hemocyte cell proliferation, respectively. We find that these transcripts are significantly lower in *Mt2*^−/−^ flies as opposed to wild type and the rescue flies (Fig. 2E), again indicating a decline in the ability of flies to mount an effective cellular transcriptional response to infection. The above data strongly suggests that Mt2 plays a key role in maintenance of healthy immune response in older flies via transcription of genes involved in the cellular arm of fly immunity. This *Mt2* function appears to become more critical as the fly ages. In the next section, we tested the transcriptional levels of gens that code for the anti-microbial peptides, *Diptericin (Dipt)*, *Attacin D (AttD)* and *Drosomycin (Drs)*. These genes are activated by Toll/NFκB or IMD/NFκB signaling and serve as readout for these pathways.

**Fig. 2:**
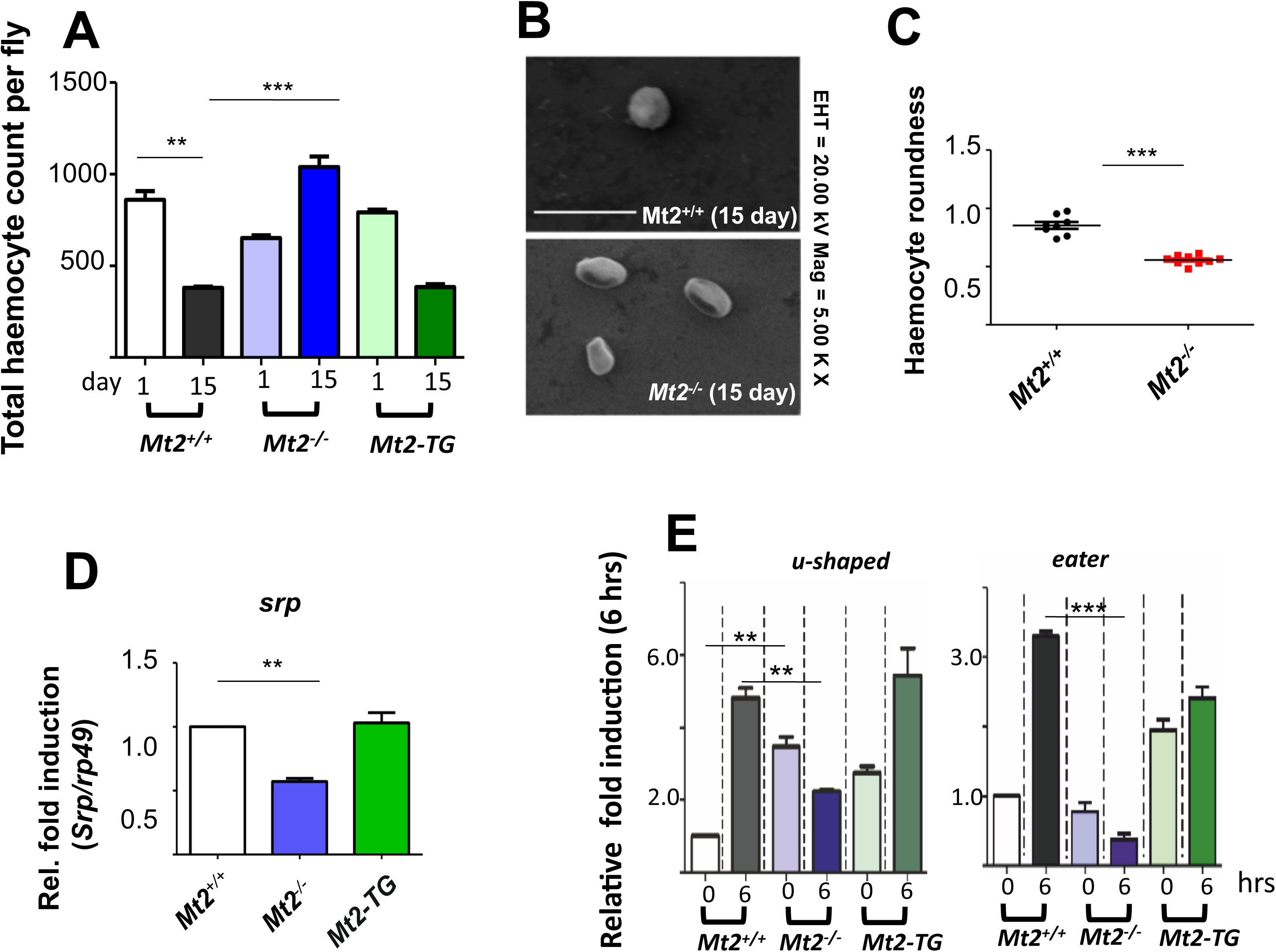
Hematopoiesis is disturbed in *Mt2*^−/−^ animals. A. The total hemocyte count for 1 day and 15 day old *Mt2*^+/+^, *Mt2*^−/−^ and *Mt2*-*TG* flies indicates increase in plasmatocyte number with age for *Mt2*^−/−^ animals, a trend opposite to that of the controls. The data shown represents three independent biological replicates with 15 males per replicate. 1Way ANOVA followed by tukey test was performed for statistical analysis. N=3, n=15.
B. Plasmatocytes from 15 day old flies imaged using SEM at 5K magnification show that *Mt2*^−/−^ show ’sickle-cell’ morphology, as compared to nearly round cells seen in wild type flies. The bar indicates a linear scale of 2 μM.
C. The linear dimensions of individual cells from SEM images were analyzed using ImageJ and the roundness for each cell was plotted in GraphPad Prism version 5. N=3; n=4. Student’s t-test was used for statistical analysis.
D. *serpent(srp)* transcript levels, as measured by real-time qPCR were reduced by half in *Mt2*^−/−^ animals, without infection. 1WAY ANOVA followed by Tukey’s test were performed as a test of significance. N=3, n=5.
E. Real time qPCR for *eater and u*-*shaped* was carried out for 15 day old flies’ pre and post infection. The data is a mean of three independent biological replicates (N=3), with 5 animals per experiment (n=5). Interestingly, the production of AMPs and cellular immunity players appear to be lowered with age in *Mt2*^−/−^ flies in comparison with *Mt2*^+/+^.

### *Mt2*-null animals show age dependent decline in AMPs

Real time PCR data was used to measure whole animal transcript levels of *Dipt*, *AttD* and *Drs* for day 2 and day 15 post eclosion. For this experiment, males of the correct age were infected with *E. coli* and transcript levels were measured at 0 and 6 Hours post-infection. Wild type flies two days post eclosion, showed 275 fold, 75 fold and 100 fold increase in transcripts for *Dipt*, *AttD* and *Drs* respectively on infection. In contrast, all three genes showed 800 fold, 140 fold and 175 fold increase in transcripts for *Dipt*, *AttD* and *Drs* respectively (Fig. 3A) for *Mt2*^−/−^. For *Mt2*-*TG*, the transcript levels were similar to *Mt2*^+/+^. This suggests, that in younger (2 Day) *Mt2*^−/−^ animals, the humoral immune response is robust and may be stronger than that in wild-type flies. For15 day old *Mt2*^−/−^ flies, transcripts of all three genes, *Dipt*, *AttD* and *Drs* were minimally responsive to infection (Fig. 3B), indicating a breakdown in signaling or lack of transcription by the NFκBs, DL, Dif and REL.

**Fig. 3:**
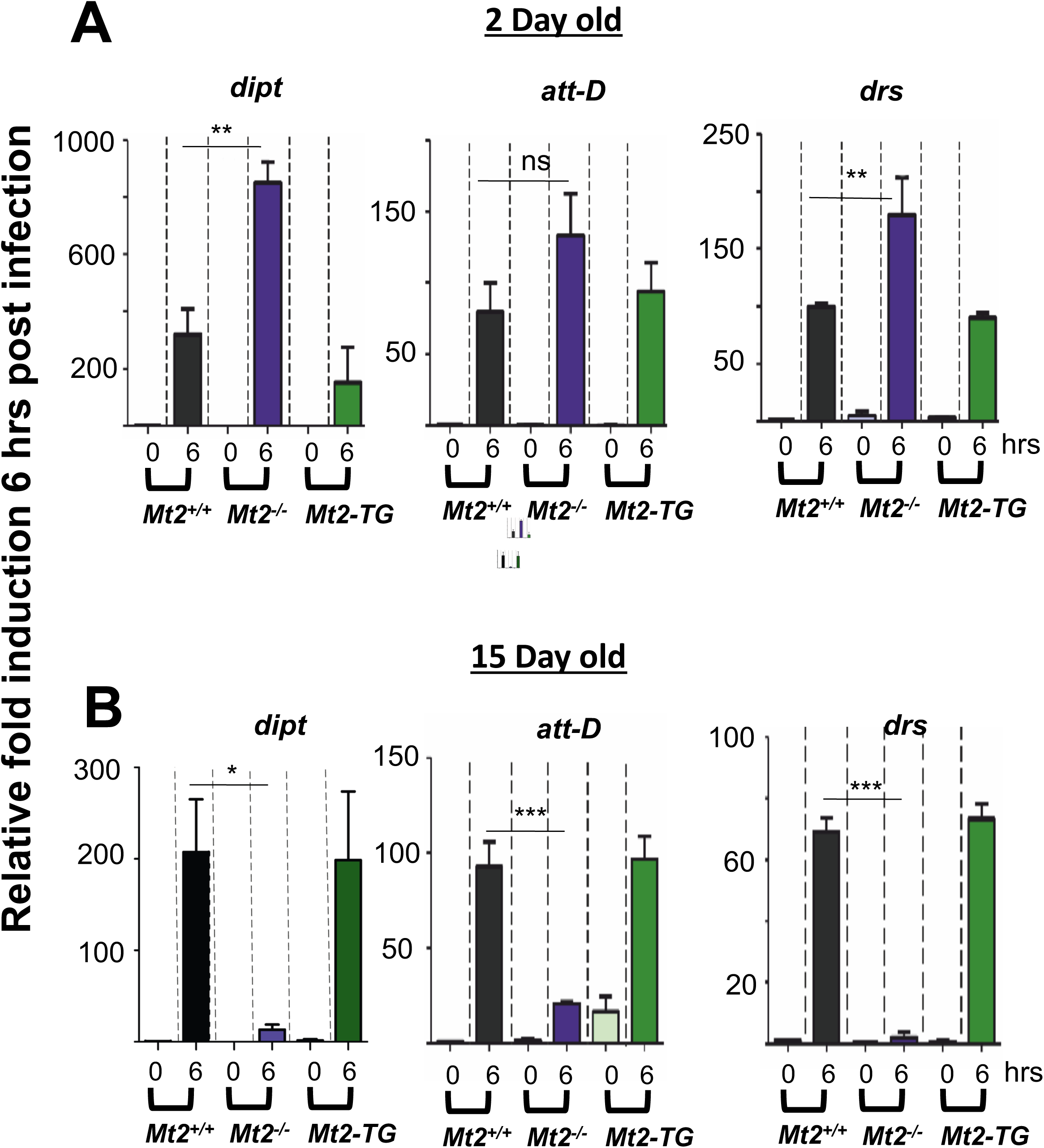
Transcriptional response by AMP genes to infection is weaker in 15-day old *Mt2*^−/−^ flies. Real Time qPCR was used to measure levels of *dipt*, *att*-*D and drs* in response to infection for 3 day and 15 day old adult *Mt2*^+/+^, *Mt2*^−/−^ and *MT2*-*TG* flies. Flies were infected and transcript levels measured at 0 and 6h post infection. Transcripts were normalized to *rp49* and relative fold values (6 hrs / 0 hrs) were plotted. 1way ANOVA followed by Tukey’s test was performed as a test of significance. N=3, n=5. A. Three day old flies show strong activation of all three AMPs. Activation of AMPs in *Mt2*^−/−^ animals is stronger, with *dipt* and *drs* levels being statistically significant.
B. Fifteen day flies show significantly lower levels of activation for all three AMPs, post infection.

### Mt2 regulates lipid homeostasis in the ageing fly

The altered shape of hemocytes at day 15 led us to profile the lipid content of *Mt2*^−/−^ animals 2-15 day post eclosion. A TAG-specific TLC analysis of the total adult fly lipidome from 2-15 days old showed significant decrease in triglycerides in *Mt2*^−/−^ animals. There appeared to be a 30% decrease in Triglyceride (TAG) levels based on quantitation of TLC bands from day 1 to day 15 (Fig. 4A). MS based quantitative lipidomics was then used to measure changes in the total lipidome for 15 day old flies (Fig. 4B; Suppl. Fig 1B). We found that the immuno-modulatory lipids, sphingosine-1-phosphate (S1P) and ceramides of varying fatty acid chain lengths, accumulated 2-3 fold in *Mt2*^−/−^ flies as compared to their WT counterparts. Concomitantly, the downstream products of sphingolipid metabolism (Fig. 5A), TAGs and phosphoethanolamine (PE) showed a ~25% decrease in *Mt2*^−/−^ flies (Suppl. Fig. 1B). The levels of lipids in the *Mt2*-*TG* rescue line was comparable to wild type. We found that several other lipid classes including neutral lipids, phospholipids (except PE), sphingomyelins, and sterols remained unchanged indicating a specific role of Mt2 in regulation of sphingolipid metabolism (Acharya and Acharya, 2005; Kraut, 2011; Saba and Hla, 2004), especially those important in immune signaling (Rivera et al., 2008). Next, we checked if, as in case of immune regulation, Mt2 also regulates lipid homeostasis in age-dependent manner. And indeed, Mt2^−/−^ showed comparable levels of S1P till day 3 post eclosion, but, by day 5, S1P starts to accumulate in these mutants as compared to controls (Fig. 4C). This accumulation is more profound as the fly ages (Fig. 4C). This accumulation of S1P led us to probe if the enzyme Sply, that converts S1P to PE (Fig. 5A), is affected. We observed a direct correlation between S1P accumulation and the failure of *Mt2*^−/−^ flies to increase Sply activity with age as compared to controls (Fig. 4D).

**Fig. 4:**
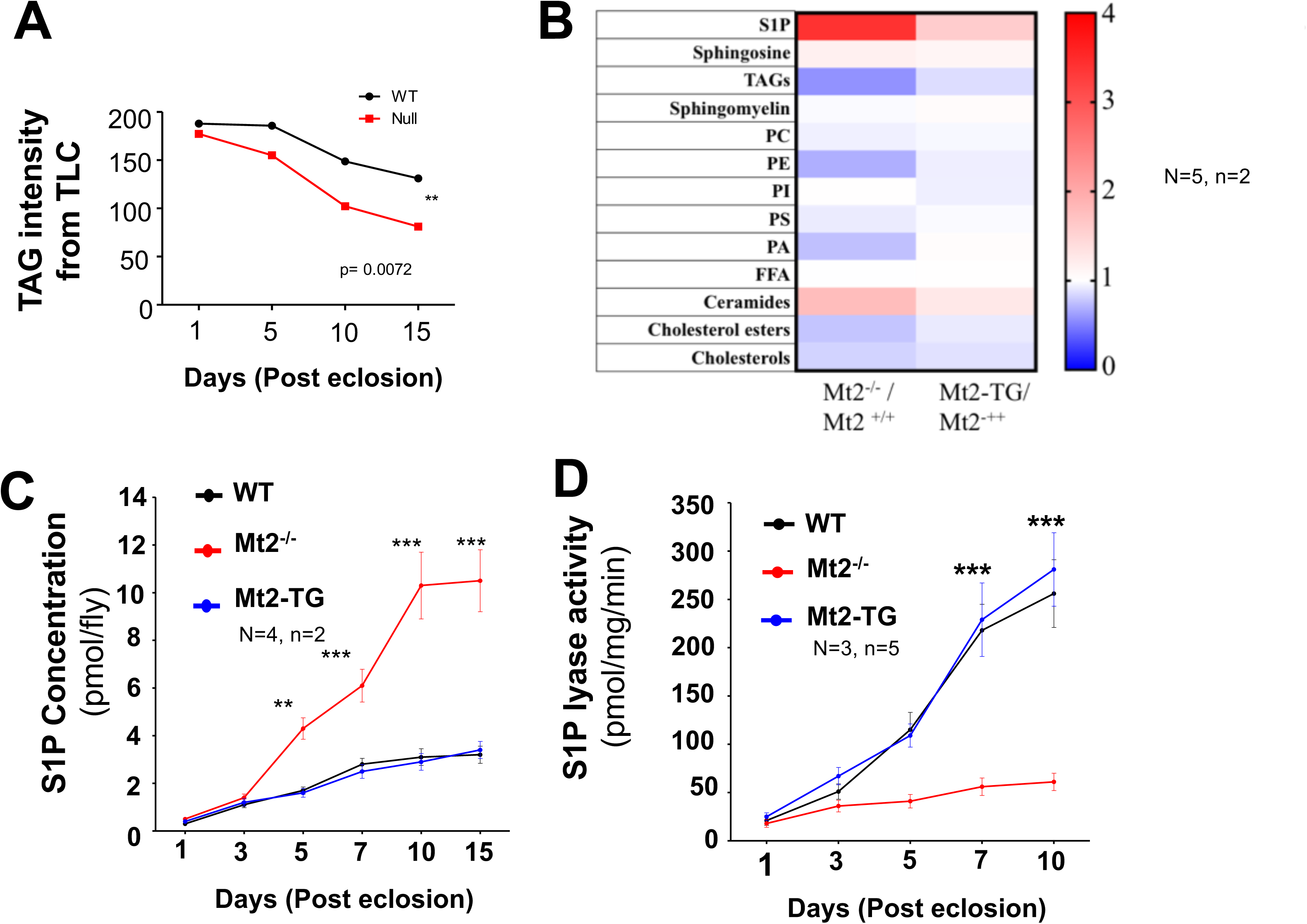
Spingosine-1-Phosphate, Ceramides levels increase while Triacylglycerol levels fall with age in *Mt2*^−/−^ flies. A. Age dependent drop in TAGs as measured by decrease in band intensities, separated by thin layer chromatography. Chi square test for trend was used for analysis. N=1, n=15.
B. Heat map that summarizes fold changes in categories of lipid moieties compared between *Mt2*^−/−^ and *Mt2*-*TG*, when normalized to Mt2^+/+^, for fifteen day old adult male flies. Red color indicates increase while blue color is fold decrease. S1P and Ceramide levels are 4 fold and 2 fold higher, respectively in *Mt2*^−/−^ flies, while Sphingomyelin, Sphingosine, Free fatty acids and overall phospholipid levels do not change significantly. TAGs, PE and PA show ~2 fold decrease in levels. Data for individual lipid moieties can be found in *Suppl. Fig. 1B* and *Suppl. Tables.* N=5, n=2.
C. Age dependent assay measures S1P levels in adult flies from day 1 to day 15. When compared to *Mt2*^+/+^ or*Mt2*-*TG*, S1P levels in *Mt2*^−/−^ flies start accumulating 5 days post eclosion. By day 10, the levels are approximately 5-fold higher than that of controls. N=4, n=2.
D. Enzyme activity assay shows change in Sply activity in Mt2^−/−^ adult whole body extracts from day 1 to day 15. Sply activity does not increase in *Mt2*^−/−^ flies with age as compared to controls. N=3, n=5.

**Fig. 5:**
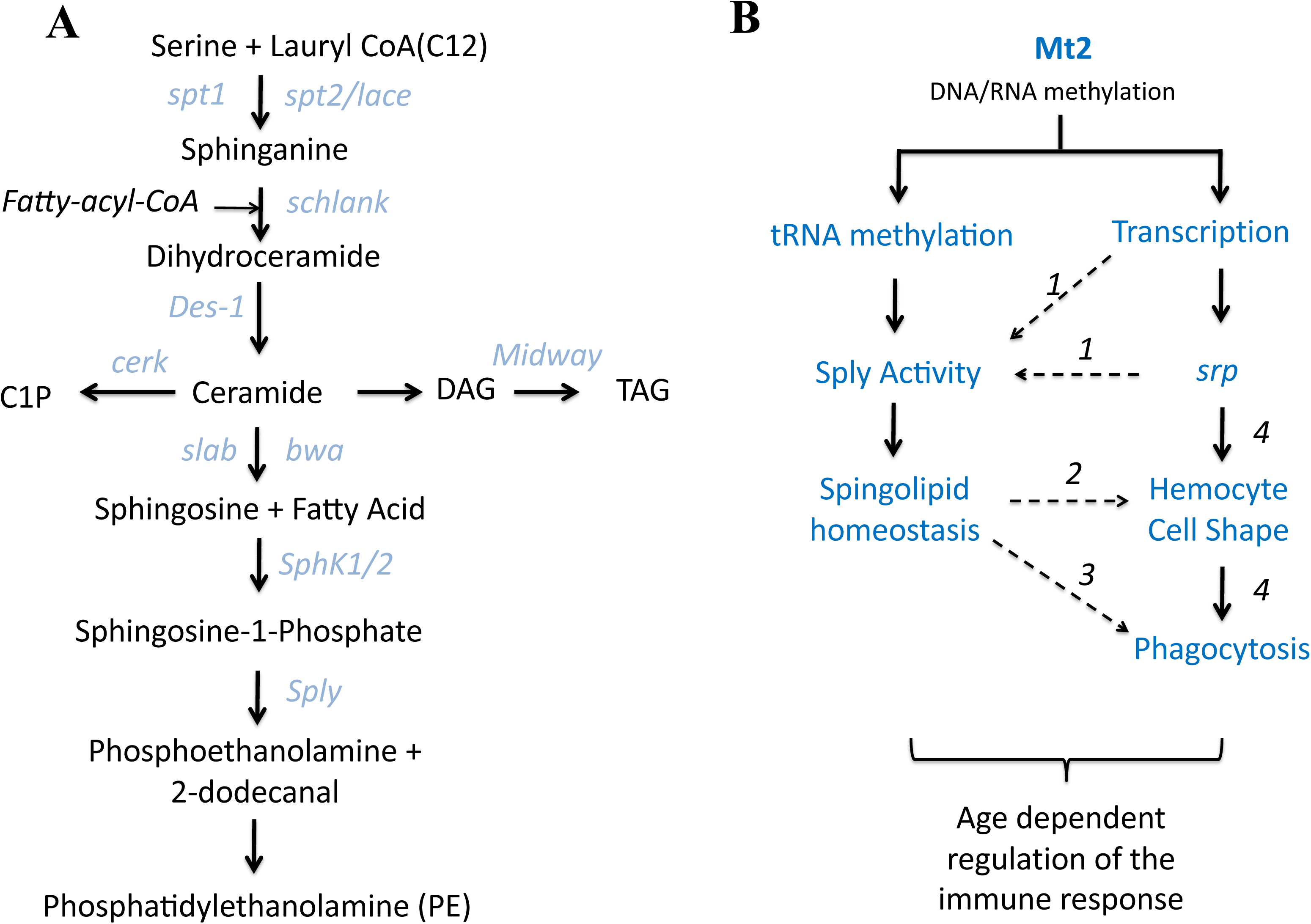
Mt2 has a systemic role in providing a robust, age dependent immune function in flies. A. Sphingolipid metabolic pathway in *Drosophila.* Metabolites are in black font while enzymes that are implicated in their conversion are in blue. Lipidomics data suggest age dependent changes in *Mt2*^−/−^ flies, with increase in S1P levels, a result in agreement with decrease in activity of Sply. Decrease in Sply activity may also explain the reduction in levels of PE. TAG levels also fall with age.
B. Model for a role for Mt2 in immunity and aging. Mt2 appears to function by regulating both the cellular and humoral arms of the innate immune response in adult flies, with lipid metabolism being a critical component for a robust response. At the molecular level, this effect would be via methylation of DNA which will regulate transcription or via methylation of tRNA, which would regulate tRNA stability and thus affect total protein activity. The model incorporates data from this study (arrows) as well as interactions found in literature (dotted arrows). The number on the dotted line indicates the source of the data. *1*(Oskouian et al., 2005), *2* (Adada et al., 2015; Kraft, 2016), *3* (Hinkovska-Galcheva et al., 2003; Tafesse et al., 2015), *4* (Ramet et al., 2002; Shlyakhover et al., 2018).

## DISCUSSION

Organisms have to manage energy in order to survive. Energy homeostasis is dependent on energy uptake, storage and expenditure. Since feeding is a discontinuous process, energy is usually stored in the form of carbohydrates, proteins or lipids to maintain a continuous supply in times of need. The *Drosophila* fat body, oenocytes, gut, malphigian tubules and special regions of the nervous system play key roles in metabolic regulation and energy homeostasis. Metabolic pathways are conserved between mammals and the fly allowing *Drosophila* to serve as a powerful model system to get a better understanding of functioning of complex metabolic networks (Owusu-Ansah and Perrimon, 2014; Padmanabha and Baker, 2014; Rajan and Perrimon, 2013; Schlegel and Stainier, 2007) including those of lipids. A finely tuned network of regulators and inter-organ communication is necessary to balance the energy intake, storage and expenditure of energy, whereby a deregulation of such networks can cause malfunction and disease.

Lipids, in addition to being storage molecules and playing structural roles in membranes, have increasingly been shown to have roles in signaling. Lipids, along with enzymes that modify and interconvert lipids constitute complex lipid signaling networks responsible for cellular and organismal homeostasis (Owusu-Ansah and Perrimon, 2014; Palm et al., 2012)(Fig. 5A summarizes *Drosophila* sphingolipid metabolic pathways). In sphingolipid metabolism levels of storage metabolites such as S1P, ceramides and TAG have to be maintained in a dynamic manner for cellular homeostasis. *Drosophila* mutants have contributed to insights into critical roles for sphingolipids in biological function. For example, mutants for sphingosine kinases (*Sphk*), which generate the important intra and intercellular signaling molecule S1P, and S1P-lyase (*Sply*) (Lovric et al., 2017), which breaks S1P down, have interesting developmental defects. *Sply* mutants show severe flight muscle defects as well as activation of apoptosis in reproductive organs (Herr et al., 2003; Phan et al., 2007), presumably by accumulating S1P. *Sphk* mutants should have reduced S1P and accumulate Sphingosine. *Sphk2* mutants, in fact, have flight defects and reduced fecundity (Herr et al., 2003). *Sply* phenotypes can be rescued by mutations in *lace*, which codes for a serine palmitoyl transferase that is a critical rate limiting step for ceramide synthesis. Ceramides act as regulators of apoptosis and are also shown to directly affect phophorylation of retinoblastoma (Rb) in response to TNFα signaling (Lee et al., 1996). S1P, in mammalian context, is shown to function via GPCRs and is suggested to regulate events such as cell shape change in PC12 cells (Edsall et al., 2001).

We find that *Mt2*^−/−^ mutants are unable to deal with infections as they age. As early as 15 days post eclosion, mutant flies are severely compromised in terms of their ability to clear infection, with plasmatocytes having disproportionately high number but defective shape. This finding parallels an imbalance in lipid homeostasis. Quantitative lipidomics confirms that S1P levels are four-fold higher than in controls, though sphingosine levels are normal. This would suggest, based on our current understanding of S1P regulation that Sply activity may be reduced. This is confirmed by enzyme activity assays in fly lysates that show reduction of Sply activity (Fig. 4D). Reduction in activity does not appear to be a result of lower transcript levels as *sply* mRNA levels do not decrease significantly (data not shown). The phenotypes could be due to errors in translation due to tRNA methylation defects earlier reported in *Mt2* mutants. Alternatively, *sply* could be regulated in tissue/immune specific manner in flies in a way similar to seen in *C. elegans*, where expression of S1P lyase is regulated by GATAA-like transcription factors and limit its expression to gut (Oskouian et al., 2005). In *Drosophila*, Srp is one of the GATAA-like transcription factors known to regulate Aldehyde dehydrogenase (Abel et al., 1993) and immune specific genes in tissue specific manner (Petersen et al., 1999; Senger et al., 2006). It would be interesting to see whether there is any regulatory link between Srp and Sply and if Mt2 plays a key role in this communication.

The lipidomics data also suggests that Ceramide levels are higher while neutral lipids are reduced suggesting more than one link in lipid metabolism affected in *Mt2*^−/−^ mutants. The three-fold increase in Ceramide levels suggest either a backflow from Sphingosine, which is maintained at normal levels, or increased activity of enzymes that metabolize Ceramide. Curiously, TAG levels are low which may suggest that the conversion of Ceramide to TAG via DAG is overactive in order to compensate for the low TAG levels. The decreased TAG levels suggest either a need for energy in the animal of a malfunction of enzymes (Fig. 5A) maintaining homeostatic levels of TAG.

The defective ‘sickle’-shaped hemocyte morphology (Fig. 2B, C) suggest architectural problems in maintaining the shape of the cell; with lipid homeostasis being a prime candidate. Since sphingholipids are critical for membrane architecture, the aberrant morphology and subsequent inability to function as macrophages may be a consequence of a reduction of sphingolipids. Mutations in S1P lyase have been implicated in regulation of cell shape with our data suggesting its malfunction being a specific cause of sickle morphology.

The correlation between imbalance in lipid homeostasis and host defense is a less explored area of research. It is understood that with environmental or nutrient stress, accumulation of lipids or signaling intermediates can interfere with immune regulation (Ertunc and Hotamisligil, 2016). Sphingolipid imbalance has been specifically linked to a number of studies (Bandhuvula and Saba, 2007; Bektas et al., 2010; Park et al., 2013; Rivera et al., 2008; Vijayan et al., 2017; Weber et al., 2009), but universal mechanisms are lacking.

Our study puts the spotlight on age-dependent regulation of lipid homeostasis and immune function. Mt2 activity, either through regulation of transcription of critical genes or by regulation of translation of protein products is important for a robust immune response in the aging animal (Fig. 5B). Absence of Mt2 function triggers an age-dependent decline in both the cellular and humoral arms of the immune response. The mechanism that Mt2 utilizes for such a systemic regulation is unclear because of the uncertainties related to Mt2 function in *Drosophila.* Mt2 function has a history of dispute (Krauss and Reuter, 2011; Schaefer and Lyko, 2010; Yoder and Bestor, 1996) over its importance in the growth and development of the organism and also its molecular function. Low levels (0.1 – 0.6%) of 5-genomic methylcytosine (5mC) have been detected in *Drosophila* (Capuano et al., 2014; Panikar et al., 2015; Takayama et al., 2014) with dynamic, developmental stage specific alteration in methylation patterns in *Mt2* null animals (Panikar, 2018; Takayama et al., 2014). Under normal conditions, complete knockdown of Mt2 has no visible survival defects, not only in flies, but also in rat and plant models (Goll et al., 2006). This led to a belief that *Mt2* is not a vital gene for the organism. We, along with others, show that Mt2 is required for increased lifespan under stress conditions. Here, in addition, we propose a novel function for Mt2 in regulating steady increase in Sply activity, a phenomenon essential to keep S1P levels in check as the fly ages. In absence of Mt2 function, this regulatory mechanism is lost, S1P starts to accumulate with age, leading to adverse effects on the ability of the fly to deal with infection. Our study, thus uncovers a novel and unexpected relationship between Mt2 mediated activity, age associated lipid homeostasis and the robust nature of the immune response.

## EXPERIMENTAL PROCEDURES

### Flystocks

Wild-type, W1118 (*Mt2*^+/+^), *Mt2* null *Mt2*^−/−^ (*Dnmt2^99^* (Schaefer et al., 2010)) and transgenic rescue *Mt2*-*TG* (w1118; pGeno>>Dnmt2-EGFP (Schaefer et al., 2008)) flies were maintained on standard corn meal medium at 25°C. *Mt2*^−/−^ and *Mt2*-*TG* flies were provided by Dr. Frank Lyko (DKFZ, Germany) and Dr. Matthias Schaefer (MFPL, Austria, Vienna) respectively. The lines were validated by measuring transcript levels in *Mt2(*^−/−^), genomic PCR to confirm deletion as described by (Schaefer et al., 2010) and PCR followed by sequencing to confirm *Mt2*-*TG* flies (data not shown).

### Survival Analysis

For survival assays, 30 three day old males from each genotype (*Mt2*^+/+^, *Mt2*^−/−^ *and Mt2*-*TG*) were maintained on standard medium at 25 °C or 29 °C. Another set of 30 flies, each pricked with 1X PBS or a 20 Hour old culture of ampicillin resistant *E. coli* (DH5α). Dead flies were removed every day and food vials were changed every day. Surviving flies were scored for two weeks at both temperatures i.e. 25 °C as well as 29 °C. Thirty flies were tested for each genotype for each condition in biological quadruplets. Kaplain-Meier and Log Rank (Mantel-Cox) test was performed using GrapPad Prism 5.0 to analyze the data.

### Bacterial Clearance Assay

2 day, 15 day and 30 day old male flies from each genotype (*Mt2*^+/+^, *Mt2*^−/−^ and *Mt2*-*TG*) were pricked with *E. coli* and kept at 25 °C for 6 hours. Four live flies from each genotype were surface sterilized using 70% ethanol. Flies were air-dried and washed twice with autoclaved MQ under sterile condition, crushed in 100μL of LB and plated on Ampicillin containing Agar plates. Colony count was taken and plotted in the form of bar graph. The experiment was repeated thrice for each genotype. Results were analyzed using One-way ANOVA in GraphPad Prism 5.0.

### Hemocyte count

Hemolymph was extracted as described (Neyen et al., 2014). In brief, 15 flies (1 day and 15 day old males) from each genotype were placed on a 10 μM filter spin column (ThermoFisher, Cat. No. 69705), covered with 4 mm glass beads (Zymoresearch, Cat. No. S1001 Rattler™) and centrifuged for 20 min at 4 °C, 10 K rpm in a microcentrifuge. The extracted hemolymph was collected in 20μL of 1X PBS solution containing 0.01% phenylthiourea, to prevent melanization of hemolymph, and counted using a Brightline hemocytometer as described (Kacsoh and Schlenke, 2012). The experiment was repeated thrice for each genotype. The total number of hemocytes per fly was plotted and One-Way ANOVA was performed in GraphPad Prism 5.0 to analyze the results.

### Counting Crystal Cells in Larvae

Crystal cells were visualized by heating thirty 3^rd^ instar larva from each genotype (*Mt2*^+/+^ and *Mt2*^−/−^) at 60 °C for 10 minutes. Photographs were taken using Zeiss microscope (AxioVision) and crystal cells were counted using ImageJ software. The results were analysed in GraphPad Prism 5.0 using Student’s t-test.

### Real time PCR

Total RNA was extracted from all the samples 0 and 6 hours of post infection (Direct-zol™ RNA MiniPrep Cat. No. R2050). cDNA was then synthesized from 1 ug total RNA using High capacity cDNA synthesis kit (Cat No. 4368814). Quantitative PCR experiments were accomplished with a StepOnePlus machine (ABI) and using SYBR Green (ABI, Catalog # 4368706). Relative gene expression was calculated after normalization to the control RpL32/rp49 mRNA. The primer sequences are available as *Suppl. Table 2.*

### SEM (Scanning Electron Microscopy)

Hemocytes from 1 and 15 day old adult males for Mt2 ^+/+^, Mt2 ^−/−^ genotypes were isolated as described in an earlier section. The drop of hemocytes was allowed to settle down on silicon wafer for 30 minutes at room temperature. Hemocytes were then washed with 20μL of 1X PBS (Phosphate buffered saline, pH 7). 20μL of fixing solution (50% ethanol, 5% acetic acid and 1% Para-formaldehyde) was added on to cells and kept overnight at 4^°^ C in a clean chamber. Next day cells were washed with 50%, 70%, 90% and 100% ethanol, air dried and imaged using Zeiss FE-SEM. Circularity index was calculated using Image J software (Circularity plugin). A perfect circle gets indicated by circularity value of 1.0 and as this value gets closer 0, it indicates an elongated polygon.

### Lipid extraction for thin layer chromatography (TLC)

Lipid isolation was done using a modified Folch extraction protocol (Kamat et al., 2015). Briefly, 5 whole adult males were crushed in 1ml DPBS in a glass vial and 1ml Methanol was added, and the mixture vortexed. Thereafter, 2ml of chloroform was added to these samples and vortexed vigorously. The sample was then centrifuged at 2800g for 5 minutes to separate the aqueous and organic phases. The organic phase (bottom) containing lipids was collected in clean glass vial. To enrich for phospholipids, the aqueous layer was acidified using 2.5% v/v formic acid, and re-extracted using 2 ml choloroform, and the two phases were separated by centrifugation at 2800g for 5 mins. The two organic phases were pooled and dried using N_2_ gas. The sample was spotted onto silica TLC plates using a glass capillary. The solvent system used was that of Wilfling *et. al.* (Wilfling et al., 2013) with minor modifications. The TLC was run using two different mobile phases sequentially. The first solvent was a mixture of n-hexane/diethyl ether/acetic acid (70:30:1). The first solvent was run halfway upto the top of the plate, after which the plate was air-dried. The plate was then run in solvent mixture of n-hexene:diethyl ether (59:1). The plate was dried and visualized by spraying with 10% (w/v) CuSO_4_ in 8% (v/v) H_3_PO_4_ followed by baking in the oven above 150^°^C for 20 mins. The plates were scanned and quantified using Image J-software.

### Quantitative lipidomics

All lipid extractions were done as described above, with small modifications (Kamat et al., 2015). Briefly, the 5 whole adult males were washed with PBS (x 3 times), and transferred into a glass vial using 1 mL PBS. 3 mL of 2:1 (vol/vol) CHCl_3_: MeOH with the internal standard mix (100 pmol of each internal standard listed in *Suppl. Table 3*) was added, and the mixture was vigorously vortexed. The two phases were separated by centrifugation at 2800g for 5 minutes. The organic phase (bottom) was removed, 50 μL of formic acid was added to acidify the aqueous homogenate (to enhance extraction of phospholipids), and CHCl_3_ was added to make up 4 mL volume. The mixture was vortexed, and separated using centrifugation described above. Both the organic extracts were pooled, and dried under a stream of N_2_. The lipidome was re-solubilized in 200 μL of 2:1 (vol/vol) CHCl_3_: MeOH, and 20 μL was used for the targeted LC-MS analysis. All the lipid species analyzed in this study were quantified using the multiple reaction monitoring (MRM) method (see *Suppl. Table 3*) on an AbSciex QTrap 4500 LC-MS with a Shimadzu Exion-LC series quaternary pump. All data was collected using the Acquisition mode of the Analyst software, and analyzed using the Quantitate mode of the same software. The LC separation was achieved using a Gemini 5U C-18 column (Phenomenex, 5 μm, 50 x 4.6 mm) coupled to a Gemini guard column (Phenomenex, 4 x 3 mm, Phenomenex security cartridge). The LC solvents were: For positive mode: buffer A: 95:5 (vol/vol) H_2_O: MeOH + 0.1% formic acid + 10 mM ammonium formate; and buffer B: 60:35:5 (vol/vol) iPrOH: MeOH: H_2_O + 0.1% formic acid + 10 mM ammonium formate, For Negative mode: buffer A: 95:5 (vol/vol) H_2_O: MeOH + 0.1% ammonium hydroxide; and buffer B: 60:35:5 (vol/vol) iPrOH: MeOH: H_2_O + 0.1% ammonium hydroxide. All the MS based lipid estimations was performed using an electrospray ion source, using the following MS parameters: ion source = turbo spray, collision gas = medium, curtain gas = 20 L/min, ion spray voltage = 4500 V, temperature = 400 °C. A typical LC-run consisted of 55 minutes, with the following solvent run sequence post injection: 0.3 ml/min 0% buffer B for 5 minutes, 0.5 ml/min 0% buffer B for 5 minutes, 0.5 ml/min linear gradient of buffer B from 0 – 100% over 25 minutes, 0.5 ml/min of 100% buffer B for 10 minutes, and reequilibration with 0.5 ml/min of 0% buffer B for 10 minutes. A detailed list of all the species targeted in this MRM study, describing the precursor parent ion mass and adduct, the product ion targeted can be found in *Supp. Table 3B.* All the endogenous lipid species were quantified by measuring the area under the curve in comparison to the respective internal standard and then normalized to the number of flies. All the data is represented as mean ± s. e. m. of 5 biological replicates per group (*Suppl. Table 3*).

### Sply activity assay

Total protein was isolated from 5 flies per replicate per genotype. 15 μg of proteome was incubated with 100 μM S1P (S9666, Sigma) in a reaction volume of 100 μL in PBS at 37^°^C with constant shaking. After 30 minutes the reaction was quenched with 350 μL of 2:1 (vol/vol) CHCl_3_: MeOH, doped with 250 pmol internal standard, cis-10-heptadecenoic acid (C17:1 FFA). The mixture was vortexed, and centrifuged at 2800 *g* for 5 minutes to separate the aqueous (top) and organic (bottom) phase. The organic phase was collected and dried under a stream of N_2_ gas, re-solubilized in 100 μL of 2:1 (vol/vol) CHCl_3_: MeOH, and subjected to LC-MS analysis. A fraction of the organic extract (~ 20 μL) was injected onto an AbSciex QTrap 4500 LC-MS with a Shimadzu Exion-LC series quaternary pump. LC separation was achieved using a Gemini 5U C-18 column (Phenomenex, 5 μm, 50 x 4.6 mm) coupled to a Gemini guard column (Phenomenex, 4 x 3 mm, Phenomenex security cartridge). The LC solvents were: buffer A: 95:5 (vol/vol) H_2_O: MeOH + 0.1% ammonium hydroxide, and buffer B: 60:35:5 (vol/vol) iPrOH: MeOH: H_2_O + 0.1% ammonium hydroxide. A typical LC run consisted of 15 minutes post-injection: 0.1 mL/min 100% buffer A from for 1.5 minutes, 0.5 mL/min linear gradient to 100% buffer B over 5 minutes, 0.5 mL/min 100% buffer B for 5.5 minutes, and equilibration with 0.5 mL/min 100% buffer A for 3 minutes. All MS analysis was performed using an electrospray ionization source in a MS1 scan negative ion mode for product formation (free fatty acid from S1P). All MS parameters were the same as those described in the MS-based lipids profiling method described above. Measuring the area under the peak, and normalizing it to the internal standard quantified the product release for the lipid substrate hydrolysis assays. The substrate hydrolysis rate was corrected by subtracting the non-enzymatic rate of hydrolysis, which was obtained by using heat-denatured proteome (15 minutes at 95 °C, followed by cooling at 4 °C for 10 mins x 3 times) as a control. All the data is represented as mean ± s. e. m. of 3 biological replicates.

## ACKNOWLEDGEMENTS

Funding for this research came from intramural and Department of Biotechnology (DBT) grants to GR. DD is supported by UGC-UPE Phase II Biotechnology [UGC-262(A)(1)] and SPPU-DRDP grant. SSK acknowledges funding from the Wellcome Trust-DBT India Alliance (grant: IA/I/15/2/502058). Shabnam Patil and Jeet Kalia are thanked for technical assistance with and access to the LC-MS facility respectively at IISER Pune, and Ajeet Singh is thanked for access to the LC-MS facility at CAMS, NCL Venture Center. VA and BK are Graduate Students supported by fellowships from CSIR, Govt. of India.

## AUTHOR CONTRIBUTIONS

VA and BK designed and carried out all of the experiments. SK designed and coordinated data related to lipid measurements. DD and GR conceived, designed and coordinated the study and drafted the manuscript.

## SUPPL. TABLES (XLS files)

**Suppl. Table 1**: *Lipid data collected for fifteen day old Mt2*^+/+^, *Mt2*^−/−^ *and Mt2*-*TG animals*.

**A** (Tab 1) LC-MS quantitation of different categories of lipids.

**B** (Tab 2). Details of the Multiple reaction monitoring transitions for the different lipids measured in the experiments.

**Suppl. Table 2**: *Primers used for RT*-*PCR*, *sequencing and validation*.

**Supplementary Figure 1.**
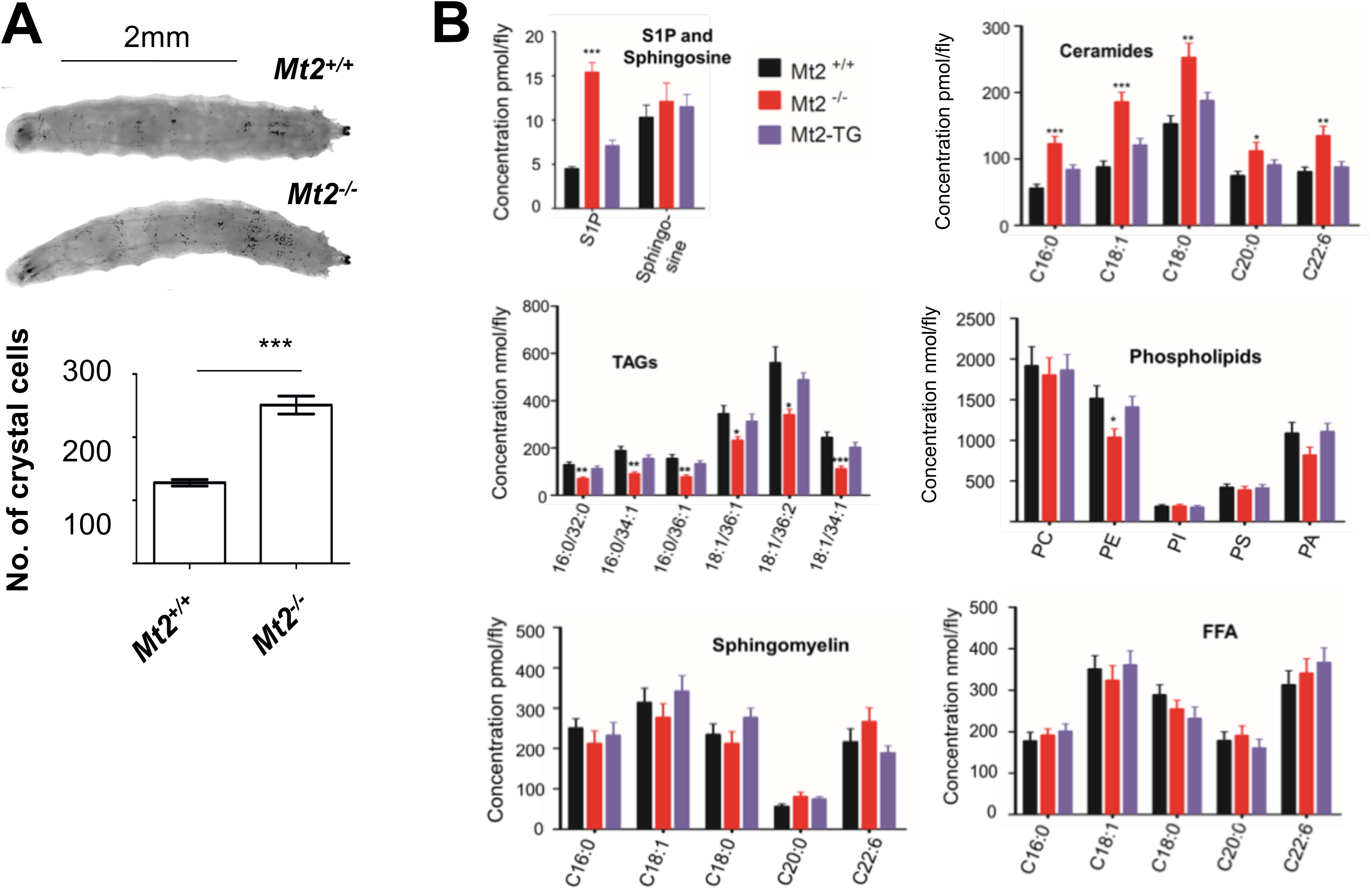
A. Crystal cell numbers are ~2-fold higher in *Mt2*^−/−^ larvae when compared to *Mt2^+/+^* suggesting a role for Mt2 in regulating larval hematopoiesis. Crystal cells were counted in three abdominal segments. N=3, n=4. This is the earliest phenotype seen in Mt2^−/−^ flies.
B. Quantitative lipidomics measuring changes in lipid moieties in 15 day old *Mt2^+/+^*, *Mt2*^−/−^ and *Mt2*-*TG* flies. Sphingomyelin, Sphingosine, Free fatty acids and overall phospholipid levels do not change significantly while S1P, Ceramides and TAGs show significant changes. 1Way ANOVA followed by tukey test was performed. N=5.

